# Highly replicated experiments studying complex genotypes using nested DNA barcodes

**DOI:** 10.1101/2025.03.18.643964

**Authors:** Molly Monge, Simone M. Giovanetti, Apoorva Ravishankar, Meru J. Sadhu

## Abstract

Many biological experiments involve studying the differences caused by genetic modifications, including genotypes composed of modifications at more than one locus. However, as the number and complexity of the genotypes increases, independently generating and tracking the necessary number of biological replicate samples becomes a major challenge. We developed a barcode-based method to track large numbers of independent replicates of combinatorial genotypes in a pooled format, enabling robust detection of subtle phenotypic differences. To construct a plasmid library of combinatorial genotypes, we utilized a nested serial cloning process to combine gene variants of interest that have associated DNA barcodes. The final plasmids each contain variants of multiple genes of interest, and a combined barcode that specifies the genotype of all the genes while also encoding a random sequence for tracking individual replicates. Sequencing of the pool of barcodes by next-generation sequencing allows the whole population to be studied in a single flask, enabling a high degree of replication even for complex genotypes. Using this approach, we tested the functionality of combinations of yeast, human, and null orthologs of the nucleotide excision repair factor I (NEF-1) complex and found that cells expressing all three yeast NEF-1 subunits had superior growth in DNA-damaging conditions. We also assessed the sensitivity of our method by simulating downsampling of barcodes across different degrees of phenotypic differentiation. Our results demonstrate the utility of NICR barcodes for high-throughput combinatorial genetic screens and provide a scalable framework for exploring complex genotype-phenotype relationships.

## Introduction

Reverse genetics experiments, in which specific genetic variants are engineered to study their phenotypic effects, are a foundational tool in biological research. They are critical for understanding gene function at multiple levels: the role of a gene can be determined through its deletion, the molecular mechanism through which it acts can be probed through targeted mutations, and its evolutionary trajectory can be explored by comparing homologous genes. Studying combinations of genetic variants further expands these insights by revealing interactions between genes or mutations. Such studies can determine whether genes have redundant functions (Nowak et al. 1997), determine the order of steps in a pathway (Avery and Wasserman 1992), uncover the molecular basis of physical interactions between or within proteins (Horovitz 1996), and elucidate how genetic variation affects phenotypic divergence and speciation (Brideau et al. 2006).

A challenge in reverse genetics experiments is the need for biological replicates. Studying multiple isogenic individuals per genotype allows researchers to distinguish the effects of genetic variation from environmental variance, experimental noise, and background genetic mutations, enabling more robust statistical conclusions. However, as experiments scale to include both combinatorial genotypes and biological replicates, the required number of samples grows rapidly. Experiments comparing larger numbers of genotypes require even more replicates per genotype to maintain statistical power, yet individually collecting data on more than a few dozen samples is often impractical. This trade-off results in researchers choosing to either limit replicates, reducing statistical power, or study only a subset of possible genetic combinations, potentially missing important genetic interactions.

Pooled experiments offer a solution to this challenge of scalability. In pooled experiments, a large population of genetically distinct individuals is created, each differing in the sequence of the genes of interest and a phenotypically inert DNA barcode to distinguish biological replicates. The pool of cells is then subjected to an assay that enriches genotypes that have a phenotype of interest, such as growth in a challenging environment or cell-sorting of a fluorescent reporter. Then, next-generation sequencing (NGS) is used to quantitatively identify the enriched genetic variants and their associated barcodes. This framework has been widely used to study the phenotypic effect of variant libraries across diverse biological contexts (Weile and Roth 2018). Here, we describe a strategy to generate and study pools of combinatorial genotypes with large numbers of replicates. In this approach, genotypic variants are successively combined together on a plasmid while generating combinatorial DNA barcodes, which we term <u>n</u>ested identification <u>c</u>ombined with replication (NICR) barcodes. We applied the NICR barcode method to generate combinations of yeast or human homologs of three proteins in the nucleotide excision repair factor I (NEF-1) complex and tracked thousands of independent replicates to examine the functionality of the combined NEF-1 complexes in yeast.

## Results

### Generation of highly replicated combinatorial genotypes

We developed a strategy to create a plasmid library of combinatorial genotypes such that each plasmid carries a barcode specifying its genotype and replicate information. For the three-subunit NEF-1 complex (Guzder et al. 1996), we sought to make all possible combinations of yeast, human, or null subunits, for 27 possible genotypes, with each genotype linked to many unique barcodes.

We started by associating our nine genetic cargos - yeast, human, or null alleles for each of the three NEF-1 subunits - to DNA barcodes that we refer to as ICR barcodes (identification combined with replication barcode; Fig. 1a). The ICR barcodes contain a 4-nucleotide hardcoded sequence specific to the cargo and a random 10-nucleotide sequence for marking independent replicates. The ICR barcode is introduced via a primer used to PCR-amplify the cargo for cloning onto a plasmid. We aimed for 500 transformed *E. coli* colonies per cargo, from which we generated nine single-gene plasmid pools (Fig. 1b).

**Figure 1.**
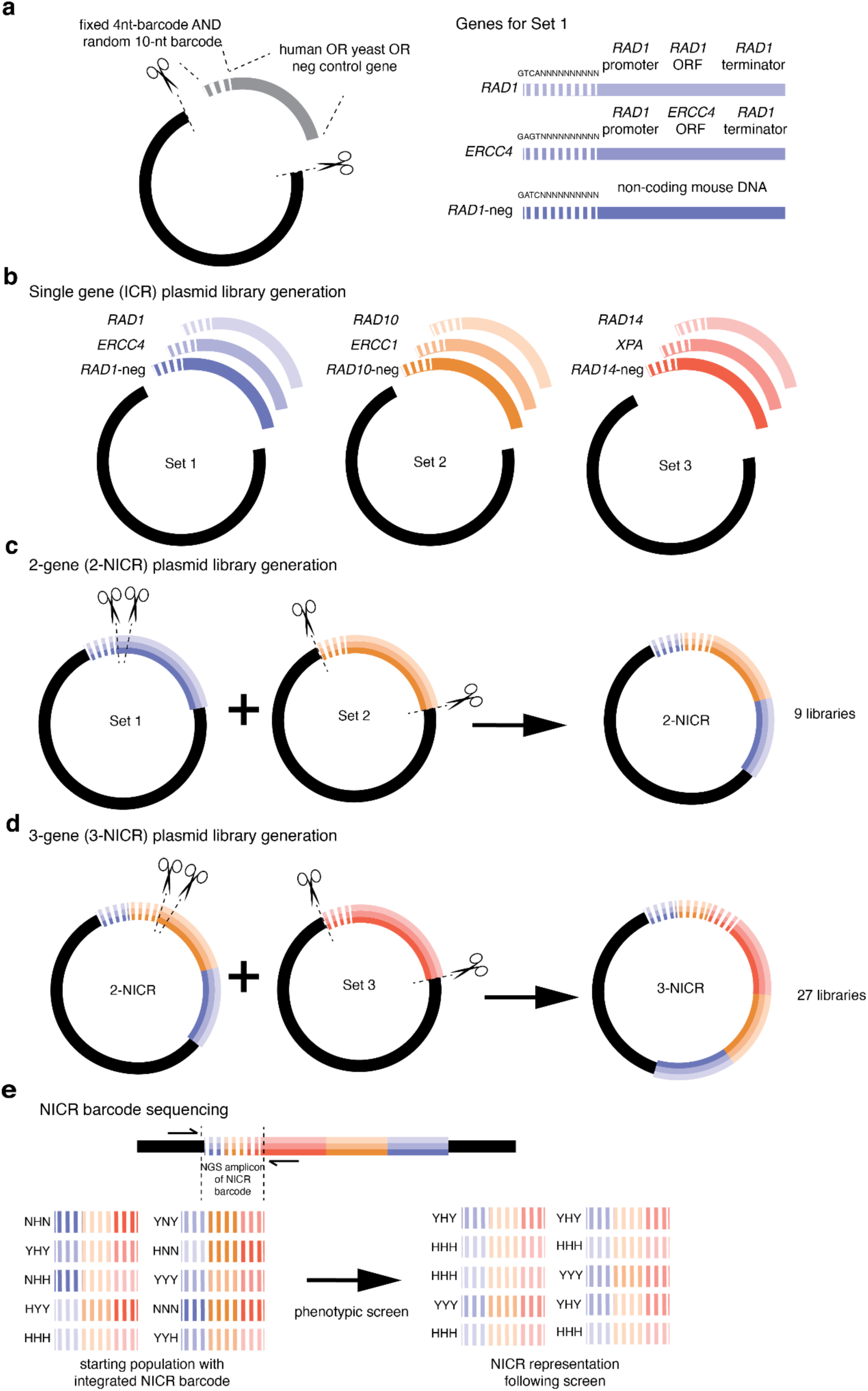
Strategy for generating highly replicated barcoded genotypes. **(a-b)** We generated nine single gene (ICR) plasmid libraries, each carrying a yeast, human, or null gene of the NEF-1 complex. Each plasmid library has a four-nucleotide genotype-specific sequence and a random 10-nucleotide sequence for tracking replicates. The yeast and human orthologs are under the control of the promoter and terminator of the yeast homolog. We combined the plasmid libraries into 27 three-gene NICR libraries. **(c)** First, the Set 2 plasmid library barcodes and cargos were cloned between the barcodes and cargos of the Set 1 plasmid libraries to generate 2-NICR libraries. **(d)** We then repeated with the Set 3 barcodes and cargos to produce 3-NICR libraries. Gene diagrams are not to scale. **(e)** Priming sequences flank the NICR-barcode for NGS sequencing of all fixed and replicate barcodes.

We then sequentially combined the single-gene libraries to create plasmids carrying all 27 possible combinations of human, yeast, or null genes for the three components of the NEF-1 complex. The nine single-gene libraries were categorized into three sets: Set 1 included libraries containing *RAD1*, its human ortholog *ERCC4*, and *RAD1*-neg; Set 2 consisted of the *RAD10*, *ERCC1*, and *RAD10*-neg libraries; and Set 3 was comprised of the *RAD14*, *XPA*, and *RAD14*-neg libraries. The sets were designed to allow restriction digestion and cloning of the barcode+gene fragment of Sets 2 and 3 to be inserted between the barcode and gene of Sets 1 and 2, respectively. Thus, in each insertion step the barcodes are placed adjacent to one another, creating larger combined barcodes that specify all the genes encoded on the plasmid. Set 1 and Set 2 were first combined to generate the 2-NICR plasmid library (Fig. 1c), which was then combined with the Set 3 library (Fig. 1d). The final 3-NICR plasmids contain a 54-nucleotide barcode that reports the identities of the three associated genes through the hardcoded components and denotes independent replicates through the random components. The final barcode is flanked by priming sites for amplification and NGS genotyping (Fig. 1e). We aimed for 500 colonies for each of the 27 combinatorial plasmid libraries.

### Highly replicated libraries contain correct genotypes and associated barcodes

We used high-fidelity long read circular consensus sequencing to perform a quality check of the contents of our initial ICR libraries by sequencing the barcodes and their adjacent genes. This allowed us to measure the rate of various cloning errors and identify barcodes associated with incorrect constructs. By creating a list of barcodes to ignore in downstream analyses, we addressed a difficulty inherent to pooled experiments, where physically removing incorrect plasmids is not feasible. Assuming the proportion of incorrect plasmids is low, the burden of carrying these plasmids through the experiment and sequencing them is minimal.

From approximately one million reads, we identified 317 to 2031 unique barcodes for each single-gene library (Fig. 2a), using a threshold of 10 reads per barcode. We analyzed the gene sequences associated with each barcode and found that the majority (68%-99%) were perfectly correct (Fig. 2a). The most common error observed was primer dimers being cloned in place of the amplified gene cargos. The maximum observed proportion of primer dimers was 31% for *RAD1*-neg, followed by 11% for *RAD1* and 10% for *RAD10*; in the remaining six plasmid pools, primer dimers accounted for less than 5% of cloned inserts.

**Figure 2.**
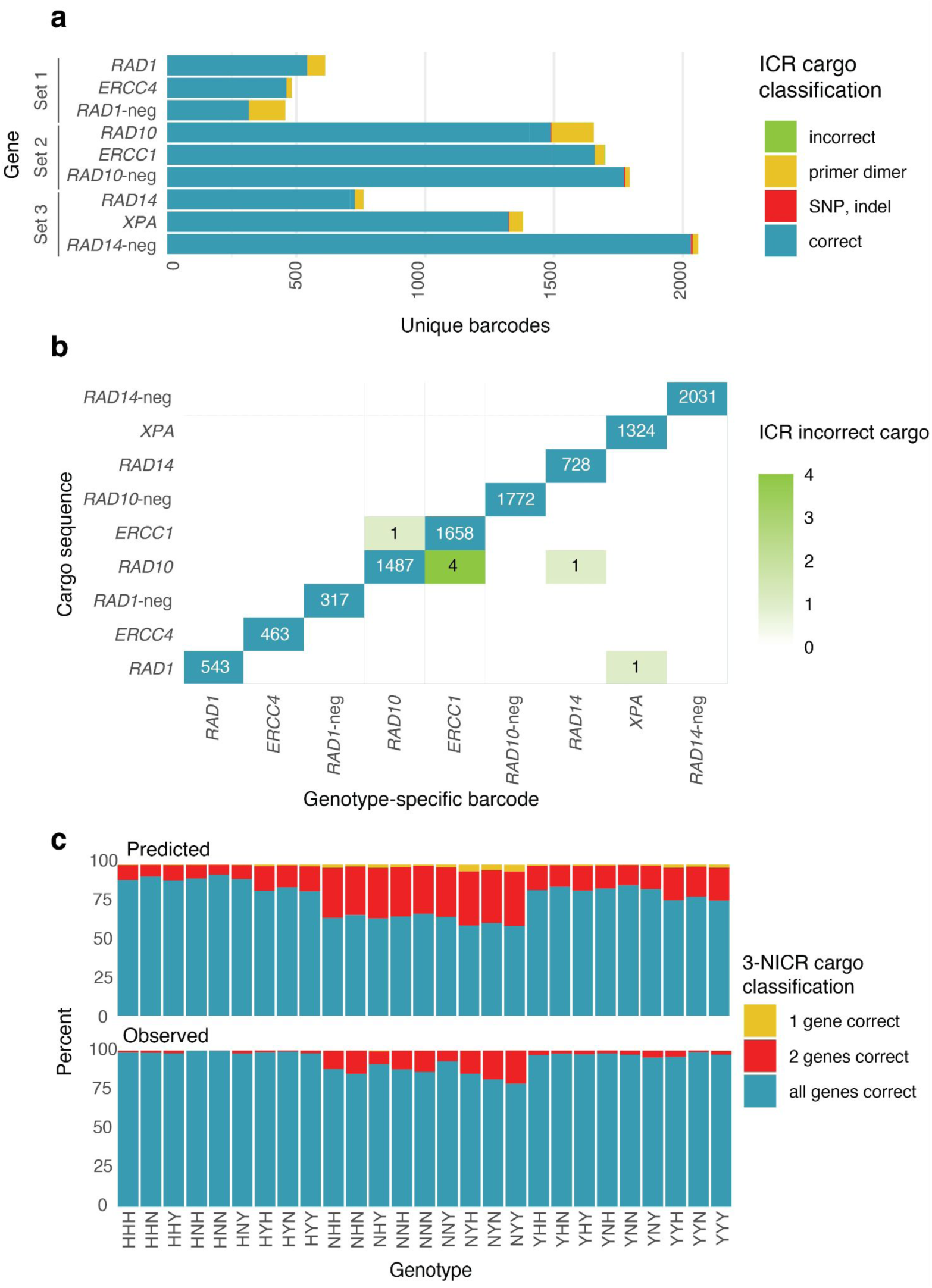
Characterization of ICR and NICR plasmid pools. **(a)** The single-gene ICR plasmids were sequenced by long read sequencing. Gene cargos were determined to be correct or incorrect. Incorrect cargos included those that contained the wrong insert (primer dimers or the wrong gene) or that had mutant versions of the right insert (SNPs, indels). **(b)** The frequency of hardcoded barcodes with the correct associated gene and with each other possible but wrong cargo. **(c)** The percent of barcodes for each 3-NICR genotype for which each 3-NICR barcode contained ICR barcodes that are associated with a correct or incorrect cargo. Each three letter code corresponds to the combination of gene cargos, in the order of Set 1, Set 2, Set 3 (Set1: *RAD1* (Y), *ERCC4* (H), or *RAD1*-neg (N); Set 2: *RAD10* (Y), *ERCC1* (H), or *RAD10*-neg (N); Set3: *RAD14* (Y), *XPA* (H), or *RAD14*-neg (N); H: human cargo, Y: yeast cargo, N: null cargo).

Spontaneous variants can arise during PCR amplification of cargos for cloning, including single-nucleotide variants or insertion/deletion (indel) variants. We found such mutations in approximately 0.5% of cargos, mostly in the form of small deletions at the very start or end of the cargo sequence (Supplementary Table 1), likely resulting from errors during synthesis of the primer used to amplify the homologs. Since the primer binding sites were chosen to be far upstream and downstream of the coding sequences, these mutations are unlikely to affect gene function, and we opted not to remove the associated barcodes from downstream analyses. Outside the primer regions, we found very few instances of SNPs and indels, affecting less than 0.05% of cargos.

Occasionally, barcodes were associated with incorrect genes, which affected less than 0.01% of barcodes (Fig. 2b). The most we observed was four cases of the *ERCC1* hardcoded barcode sequence linked to *RAD10* gene cargo. Despite separately cloning all nine single-gene libraries, this incorrect association could result from barcode primer cross-contamination, template DNA cross-contamination, or template-switching during long-read sequencing.

Next, we investigated the proportion of barcodes associated with cargos with errors in the final three-gene plasmid pools. Primer dimers were the largest population of erroneous cargos in the ICR pools. To minimize their presence, we used gel-electrophoresis-based size selection during NICR library cloning. We assessed the effectiveness of primer dimer removal by sequencing the 3-NICR barcodes using short-read sequencing (Supplementary Fig. 1) and found that their abundance was substantially reduced (Fig. 2c). Among all constructs, *RAD1*-neg had the largest proportion of primer dimer-associated 3-NICR barcodes, likely because it was always part of the “vector” during the nested cloning (Fig. 1c,d), causing its correct cargos and primer dimer cargos to migrate similarly on a gel. Still, the representation of primer dimer barcodes for *RAD1*-neg was lower in the 3-NICR pools than predicted from their representation in the ICR pool. Overall, the 3-NICR plasmid pools mostly contained barcodes associated with correct cargos (Fig. 2c); for 96% of 3-NICR barcodes, all three component ICR barcodes corresponded to correct cargos.

### Testing chimeric NEF-1 complexes in yeast

Next, we tested whether the highly replicated plasmid pools could be used to distinguish phenotypes of chimeric NEF-1 complexes in a pooled experiment. The nucleotide excision repair pathway is required to repair bulky lesions in DNA (Schärer 2013). Following bulky lesion recognition, Rad14 recruits Rad1 and Rad10 to excise the damaged DNA strand, which is then replaced through DNA synthesis. Nucleotide excision repair is required for yeast survival of the genotoxic drug methyl methanesulfonate (MMS) (Prakash and Prakash 1977). Past work has shown that the function of Rad1 and Rad10 in MMS survival can be partially replaced by their human homologs, ERCC4 (also known as XPF) and ERCC1, either individually or together (Hamza et al. 2020), while the ability to replace Rad14 with XPA has not been previously tested. To test the functionality of all combinations of yeast or human homologs of the three NEF-1 subunits, we integrated the 27 3-NICR plasmid pools into the *ura3*Δ locus in *rad1*Δ *rad10*Δ *rad14*Δ yeast, pooled all transformants together, and grew them in media containing 0, 0.005%, 0.01%, 0.015%, or 0.02% MMS. To understand the effect of MMS on each genotype’s growth, we collected cells every 12 hours for 48 hours and then sequenced the 3-NICR barcode by NGS to quantify abundance of each genotype over time.

In 0.005%, 0.01%, and 0.015% MMS, cells with the yeast homologs of all three NEF-1 genes had significantly superior growth to cells with any other genotype (Fig. 3; Supplementary Table 2). The superior growth was apparent at 24, 36, and 48 hours in 0.005% and 0.01% MMS, while in 0.015% MMS it took 48 hours to appear, likely because in 0.015% MMS growth was generally poor (Supplementary Fig. 2). Growth was even poorer in 0.02% MMS, and we did not detect differences in growth at any time point.

**Figure 3.**
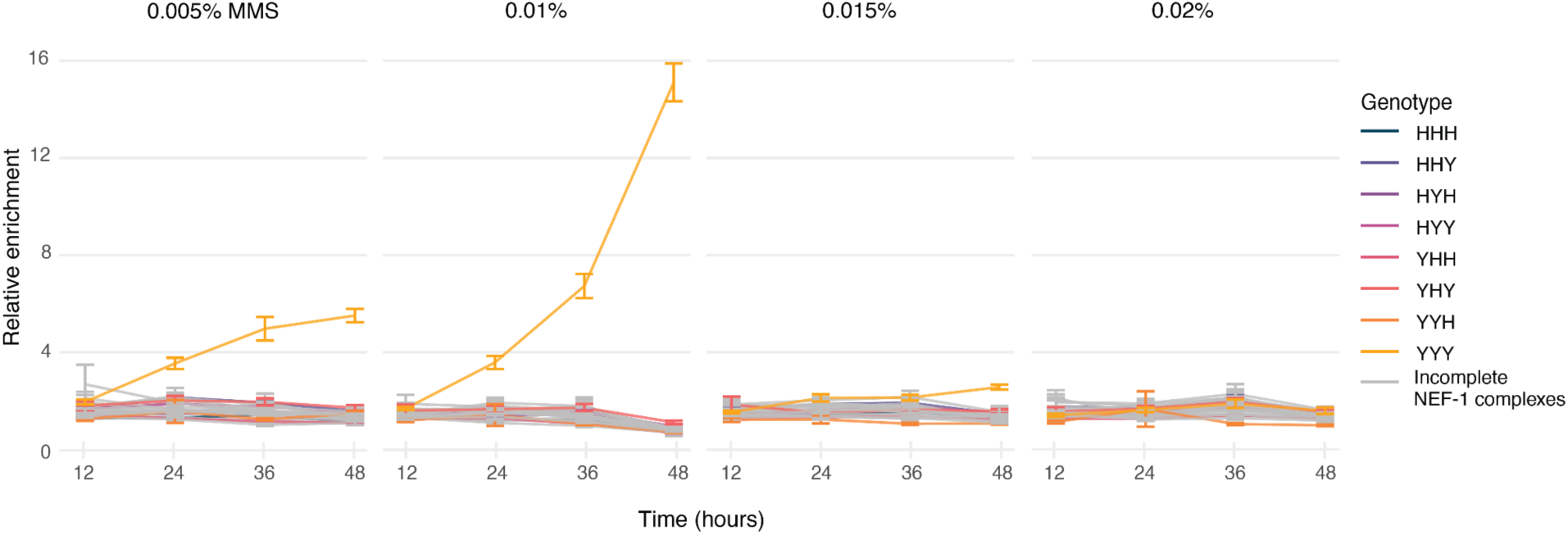
3-NICR yeast library growth in MMS. 3-NICR libraries were grown in media containing 0.005%, 0.01%, 0.015%, and 0.02% MMS. Samples were collected every 12 hours for 48 hours, and the abundance of each 3-NICR barcode was measured by NGS as a proxy for cell growth. Each 3-NICR barcode abundance was normalized to the total read count in that sample and then normalized to its abundance in media without MMS from the same time point. Each line represents the average normalized abundance across all replicates for a single genotype. Genotype identities are specified as in the three-letter code used in Fig. 2c. Error bars signify the standard error of the mean.

Surprisingly, all of the genotypes composed of a mix of yeast and human subunits had poor growth, comparable to the genotypes lacking NEF-1 subunits, whereas a previous study found that the human homologs of *RAD1* and *RAD10* could partially confer MMS resistance. The difference in results could be caused by variation in the background strain used to perform the experiments, or by our expression of the genes from the *URA3* locus.

Alternatively, it could be caused by differences in the experimental setup, such as the source of MMS used, or in the co-culture of genotypes in our study as compared to separate growth in the original study.

### Replication barcodes allow detection of small effects

Since we used random DNA barcodes generated during cloning to track biological replicates, the number of replicates scaled to the number of colonies resulting from a plasmid cloning reaction, which allowed us to generate hundreds of replicate samples. Next, we sought to determine the minimum number of replicates that would have been needed to detect the observed phenotypic differences. We randomly selected subsets of 3-NICR barcodes from the 0.005%, 0.01%, and 0.015% MMS data and determined the frequency with which we were able to detect a growth difference between strains with yeast NEF-1 compared to fully null NEF-1 (Fig. 4). In 0.01% MMS, in which yeast NEF-1 conferred the largest phenotypic benefit, we had over 90% power to detect this difference with four barcodes per genotype, while in 0.005% MMS and 0.015% MMS approximately 25 and 140 barcodes were required, respectively. These numbers of barcodes are all accessible for a pooled experiment, whereas for a non-pooled experiment the number of replicates would likely limit detection to the strongest phenotype. These results demonstrate the power of NICR barcoding to fully capture genotype-phenotype relationships with high statistical confidence by leveraging the scalability and efficiency of pooled experiments.

**Figure 4.**
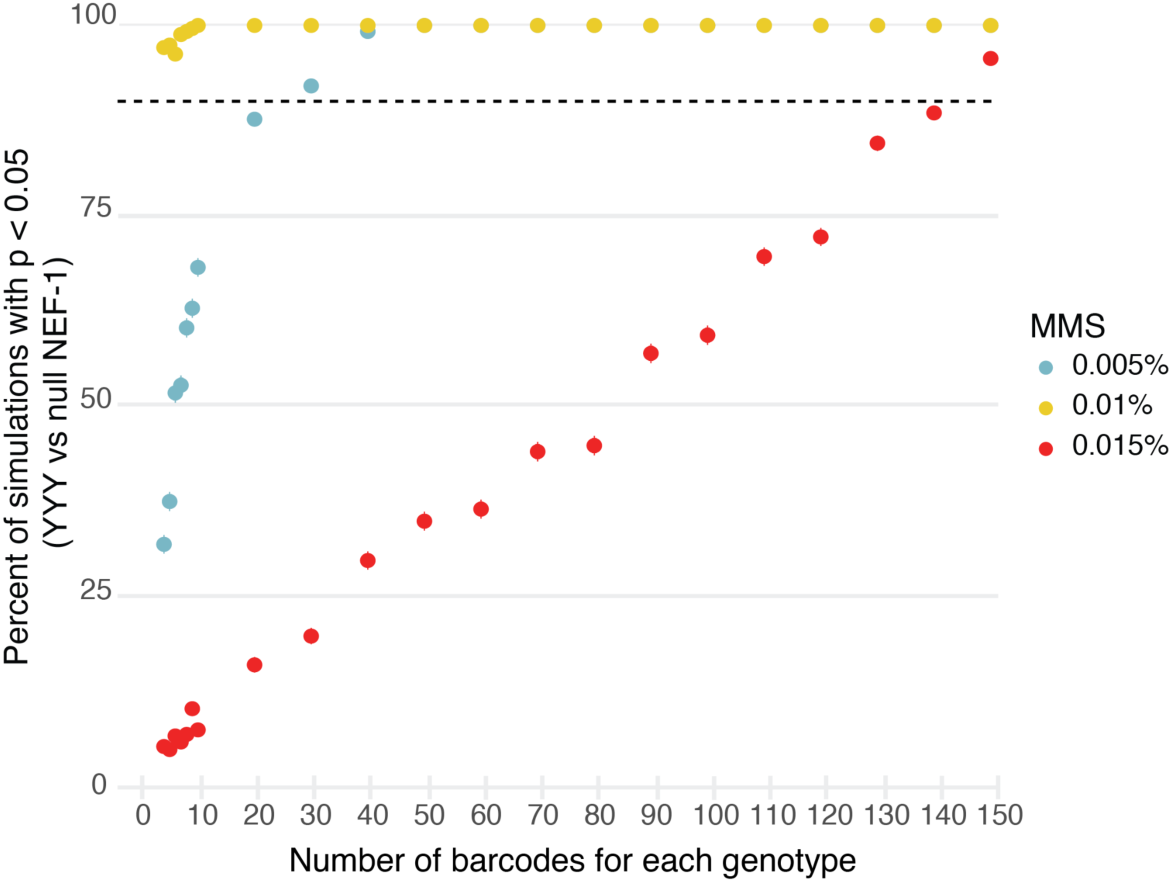
Downsampling analysis of 3-NICR barcodes. For each downsample (ranging from 4-150 3-NICR barcodes per genotype), unique 3-NICR barcodes were randomly sampled and their abundance in MMS media relative to media without MMS at the final time point was compared. We repeated the analysis 500 times for each MMS concentration. We determined the frequency that the yeast NEF-1 genotype had a statistically significant improvement in growth compared to fully null NEF-1. Statistically significant improved growth was defined as the adjusted p-value > 0.05 and effect size > 0.2. Error bars signify the 95% confidence interval.

## Discussion

Replication is essential for statistical confidence in reverse genetics experiments. However, designing an experiment with replication presents two major challenges, both of which are heightened in experiments studying genotype combinations. The first challenge is that replication can result in a large number of total samples. This often leads researchers to either choose a subset of genotypes for comparison or use a limited number of replicates. Even with these adjustments, the number of samples can be burdensome. For instance, if we had individually generated the eight human-yeast chimeric NEF-1 genotypes plus a negative control, each with four replicates, we would still have 36 samples to carry through five concentrations of MMS. This would be challenging experimentally while providing limited statistical power. Generating and studying samples in pools alleviates this difficulty. In our approach, replicates are generated as distinct colonies during a cloning reaction.

Instead of selecting individual colonies, all the colonies are collected together, allowing a single cloning reaction to generate dozens to hundreds of replicates. Then, the phenotypic assay is done simultaneously on the entire pool of replicates, which means that increasing the number of replicates increases the scale of the assay rather than the number of assays. Finally, DNA barcodes can easily be designed to distinguish millions of samples (Johnson et al. 2023), and a single NGS run can generate billions of reads for phenotypic determination.

The second challenge is to generate replicates in a way that avoids the replicates sharing background mutations that are absent from the other genotypes. If a single mutant cell line is generated and then split to produce replicates, any background mutation present in the original cell line would be shared among all replicates. The background mutations could cause phenotypic effects that would be mistakenly attributed to the targeted genetic modification rather than to these unintended mutations. Such background effects could arise from point mutations, aneuploidies, copy number variation, or even non-chromosomal heritable effects. Avoiding shared background is particularly difficult when generating combinatorial genotypes: if a genotype is generated in multiple sequential steps, such as knocking out several genes, then, to avoid shared background mutations, all replicates of all genotypes would need to be independently generated from the first step. This imposes a substantial experimental burden even with a modest number of replicates, and becomes even more demanding when aiming for high replication to enhance statistical power. Moreover, producing replicates independently from the first stage of genotype construction increases the likelihood that each replicate will acquire unique background mutations, leading to increased phenotypic variation among the replicates and decreased power to detect effects resulting from the constructed genotype.

A solution to this paradox is to have all the lines across all genotypes and replicates share a common genetic background as late as possible during genotype construction. In the approach described here, we transformed a single receiver yeast strain with a library of different genotypes delivered by integrating plasmids. Thus, most background mutations in the receiver strain will be constant among all the genotypes, while even de novo background mutations present heterogeneously in the receiver strain culture will be randomly distributed among the genotypes. We also minimized the possibility of local background mutations carried on the NICR plasmids by sequence-verifying cargos by long-read sequencing.

Pooled phenotyping requires the abundance of cells in the final pool to reflect their phenotype. This is straightforward when the phenotype has an effect on fitness, as fitter strains will grow faster than less fit strains and thereby have higher representation in the pool. In addition, assays have been developed for measuring many other phenotypes through a cellular abundance readout. Many protein-based phenotypes, including protein expression, enzymatic activity, or protein-protein interaction, can be quantitatively assayed in a pool using approaches such as fluorescence-activated cell sorting to convert non-fitness phenotypes to differences in cellular abundance (Adams et al. 2016; Matreyek et al. 2018; Amorosi et al. 2021). Single-cell sequencing has enabled genome-wide phenotypes to be measured for distinct genotypes in a pool, such as transcriptome or chromatin accessibility perturbations (Adamson et al. 2016; Rubin et al. 2019). Thus, a large diversity of phenotypes can be studied with NICR barcodes. Furthermore, the NICR barcode approach can be applied across a variety of organismal systems beyond yeast; it can be applied to any system in which DNA can be introduced and individual phenotypes measured in a pool. We envision it would be particularly useful for cell culture experiments, as heterogeneity in cell lines is common (Zhu et al. 2023). The NICR approach can even be utilized even beyond cell-based assays, as strategies have been developed to carry out pooled genetic experiments in multicellular organisms such as *Caenorhabditis elegans* and zebrafish (Parvez et al. 2021; Stevenson et al. 2023).

## Materials and Methods

### Cloning Strategy

Strains, plasmids, and oligonucleotides used in this study are listed in Supplementary Tables 3, 4, and 5.

We generated 27 plasmid pools, each carrying three gene cargos, in a three-step cloning process. Each step incorporates an additional gene to create the three-gene complex and expands the NICR barcode to account for this addition.

First, we PCR-amplified the three yeast NEF-1 genes (*RAD1*, *RAD10*, and *RAD14*), three human NEF-1 genes (*ERCC4*, *ERCC1*, and *XPA*), and three non-coding mouse DNA sequences (*RAD1*-neg, *RAD10*-neg, and *RAD14*-neg) using Herculase II Fusion DNA Polymerase (Agilent) to obtain our gene cargos. The yeast genes were amplified from yeast genomic DNA (strains MSY116, MSY119, and MSY121, respectively), and the amplicons were designed to include 228-375 base pairs (bp) of upstream promoter sequence and 299-763 bp of downstream terminator sequence, based on RNA-seq results identifying the start and end of the transcripts (Albert et al. 2014). The human genes were amplified from yeast strains that had individual human genes integrated at the homologous yeast loci (strains MSY121, MSY116, and MSY119, respectively); the integrated *ERCC4* and *ERCC1* cDNA sequences were derived from plasmids gifted from Phil Hieter, and the *XPA* cDNA sequence was ordered from the hORFeome1.1 collection (Horizon Discovery). We used mouse non-coding DNA for the negative control sequences to minimize homology to the yeast genome. We chose the sequences by scanning intronic regions of the mouse genome for neutral, noncoding DNA whose lengths were comparable to the corresponding NEF-1 gene. We used 3752 bp of a *Tyr* intron for *RAD1-*neg, 1586 bp of a *Dct* intron for *RAD10*-neg, and 1549 bp of the *Tyr* promoter for *RAD14-neg*. C57BL/6J mouse genomic DNA was a gift from Bill Pavan. Each forward primer included a 4-nucleotide hardcoded barcode that was specific to the cargo and a 10-nucleotide stretch of random nucleotides to generate the replicate barcode. Primer sequences are listed in Table 3.

The nine ICR plasmid pools were constructed by cloning these amplicons into pYTK096 (Lee et al. 2015) digested with BsaI-HFv2 and MluI. BsaI generated a linear vector fragment, and MluI cut between the two BsaI sites to further reduce undigested plasmid background. The PCR primers used to amplify the cargo insert DNA molecules introduced 20 bp of homology to the ends of the BsaI-cut vector. We assembled our nine inserts into digested pYTK096 using Gibson Assembly (NEB) and plated individually on LB+Kanamycin plates. Colonies from all nine single-gene ICR libraries were scraped and plasmid DNA was isolated using a Plasmid Plus Maxi kit (QIAGEN).

Next, we assembled our 2-NICR plasmids by combining the first and second genes of NEF-1 in our second round of cloning. Set 1 (*RAD1*, *ERCC4*, and *RAD1*-neg) plasmid pools were digested with AgeI-HF and BstEII-HF (NEB) sequentially and treated with Quick CIP phosphatase (NEB) to generate vector fragments. Set 2 (*RAD10*, *ERCC1*, and *RAD10*-neg) plasmid pools were also digested with AgeI-HF and BstEII-HF to generate insert DNA pools. We separately assembled all nine possible combinations of fragments using T4 ligase (NEB) and plated on LB+Kanamycin plates. All colonies were scraped and plasmid DNA was isolated using the Plasmid Plus Maxi kit.

We assembled our complete 3-NICR plasmids in our third round of cloning. The 2-NICR (*RAD1*/*RAD10*, *RAD1/ERCC1*, *RAD1/RAD10*-neg, *ERCC4/RAD10*, *ERCC4/ERCC1*, *ERCC4/RAD10*-neg, *RAD1*-neg/*RAD10*, *RAD1*-neg/*ERCC1*, *RAD1*-neg/*RAD10*-neg) plasmid pools were digested with NheI-HF and SphI-HF (NEB) sequentially and treated with Quick CIP phosphatase to generate the vector fragments. Plasmid pools were also digested with BsiWI-HF (NEB) to remove background vectors from previous rounds of cloning (i.e. ICR plasmids that were not digested or assembled correctly when generating 2-NICR plasmids), as in the “kill cut” strategy described by Poelwijk, Socolich, and Ranganathan (Poelwijk et al. 2019). Set 3 (*RAD14*, *XPA*, and *RAD14*-neg) plasmids were digested with NheI-HF and SphI-HF to generate the insert DNA pools. We separately assembled all 27 possible combinations of fragments using T4 ligase and plating on LB+Kanamycin plates. All colonies were scraped and plasmid DNA was isolated using the Plasmid Plus Maxi kit.

### PacBio long read sequencing

DNA was isolated from each of our initial nine ICR plasmid libraries for long read sequencing to determine whether the hardcoded barcode component was correctly paired with the correct gene cargo. ICR plasmid pool DNA was digested with NotI-HF (NEB) to excise linear fragments carrying the barcodes and gene cargos, from which 2.3 µg of DNA was cleaned twice with Ampure PB beads(Pacific Biosciences). A bead:DNA ratio of 0.45 was used to remove fragments shorter than 3 kilobases. A library was generated using SMRTbell prep kit 3.0 (Pacific Biosciences) through the adapter-barcoded workflow, and sequenced on a Sequel IIe instrument using a 8M SMRT cell with version 2.0 chemistry (Pacific Biosciences). We used a 2-hour pre-extension and a 15-hour movie. From the resultant CCS reads we used custom Python scripts to extract the 14-nucleotide ICR barcode and the associated cargo. For each unique identified barcode, the sequence of the associated cargo was classified as correct (expected error-free gene cargo matching the hardcoded barcode), misassociated (unexpected gene cargo), containing a primer dimer (short primer dimer as cargo), or containing SNPs or indels (expected gene cargo but without perfect matching). Barcodes that were associated with cargos that were both correct and incorrect cargos were discarded. We calculated the total number of barcodes for each of the 9 genotypes and the percentage of barcodes corresponding to each class. We generated a list of correct ICR barcodes for use in downstream analyses.

### Illumina sequencing

We used Illumina short read sequencing to characterize our initial 3-NICR library and to measure barcode abundances during the MMS experiment. From either a plasmid prep or yeast genomic DNA, we first amplified the 3-NICR barcode using KAPA HiFi HotStart (Roche) and custom primers. We purified the PCR product using the MinElute PCR Purification Kit (QIAGEN). We next amplified 10 ng of amplicon DNA with Nextera adapter Index primers (Illumina) and KAPA HiFi HotStart. We ran the final PCR product on a 2% SizeSelect Egel (Invitrogen) and collected the correctly sized band. Finished libraries were quantified by Qubit 4 fluorometer (Invitrogen). Libraries were diluted to 4 nM and pooled before sequencing on a NextSeq 2000 instrument (Illumina).

Illumina reads were paired using PEAR (Zhang et al. 2014) and then, using custom Python scripts, we extracted the entire 3-NICR barcode from each sequence read. From the sequencing output of the plasmid pool, we filtered for 3-NICR barcodes that contained three previously identified ICR barcodes to generate a list of correct barcodes for downstream analyses.

### Yeast Strain Construction

*rad1Δrad10Δ* (MSY101) was a gift from Phil Hieter. We constructed *rad1*Δ *rad10*Δ *rad14*Δ by replacing yeast *RAD14* in the *rad1*Δ *rad10*Δ strain via homology-directed repair with the NatMX cassette. We amplified NatMX from MSp2 via PCR using primers oMM15 and oMM16 and transformed it into the *rad1*Δ *rad10*Δ strain using a standard lithium acetate protocol (Becker and Lundblad 2001). *RAD14* deletion was confirmed via PCR. We designated the *rad1*Δ *rad10*Δ *rad14*Δ strain MSY125. We next transformed our triple knockout strain with plasmids carrying Gal-Cas9 and a gRNA to target the *ura3*Δ locus to increase the transformation efficiency of our recombinant NEF-1 cargos, creating strain MSY169.

### Yeast Transformation

We digested our 27 plasmid pools with NotI-HF, MluI-HF, BsiWI-HF, and SalI-HF and directly transformed this reaction into the *rad1Δrad10Δrad14Δ* strain (MSY169) using the standard lithium acetate transformation protocol. Digestion by NotI-HF generated linear DNA fragments consisting of the 3-NICR barcode, three gene cargos, and the *URA3* promoter, gene, and terminator, with homology arms targeting integration at the *ura3*Δ locus. Any contaminating background plasmids that did not receive one or more of the three components contained “kill cut” restriction sites (MluI-HF, BsiWI-HF, or SalI-HF) between the left and right homology arms (Poelwijk et al. 2019), such that cleavage would block genomic integration of these fragments; all three restriction sites were absent from the fully assembled 3-cargo plasmids. A single colony of the *rad1Δ rad10Δ rad14Δ* strain was grown to an optical density of 0.6 in 500 mL of media, which was used for all 27 transformations, such that any background mutations would either be fixed across genotypes or occur randomly without association to any particular plasmid genotype. Approximately 180 million cells were used for each transformation with 3.5 µg of one of the digested 3-NICR plasmid pools. We selected transformants on CSM -His -Leu -Ura (Sunrise Science Products) + 2% galactose plates, then washed resultant colonies off into 27 individual yeast transformant pools, which were frozen down as glycerol stocks.

### Testing MMS sensitivity

We made individual overnight cultures of the 27 yeast pools in 5 mL of CSM -His -Leu -Ura media. The next day, we pooled together an equal number of cells from each overnight culture and used this combined pool to inoculate 300 mL cultures that had 0, 0.005%, 0.01%, 0.015%, or 0.02% MMS to an optical density of 0.025. The cultures were grown at 30C for 48 hours. Every 12 hours we froze down approximately 60 million cells and diluted back to an optical density of 0.025 if they had reached an optical density greater than 0.2. Cells were also frozen immediately after forming the pool. We extracted DNA from the frozen cells using the yeast genomic DNA protocol with the DNeasy Blood and Tissue Kit (QIAGEN), from which we prepared Illumina libraries of the 3-NICR barcodes as above.

### Data Analysis

Using custom Python and R scripts, we used our list of filtered correct 3-NICR barcodes to track replicates and genotypes across the MMS experiment. For each barcode we normalized reads to account for differences in sequencing depth across time points and MMS concentrations. First, we calculated the percentage of reads at a given time point and MMS concentration for each correct 3-NICR barcode. To adjust for differences in baseline growth, we then normalized each barcode’s frequency to its frequency at the same time point but without MMS. A linear model was fit to the normalized barcode counts across the genotypes at a given time and MMS concentration, and genotypes significantly different from each other were detected using estimated marginal means, as implemented in the R package emmeans (v.1.10.7). Adjusted p-values were calculated using the Benjamini-Hochberg false discovery rate (FDR) correction method across all time points and MMS concentrations. Significant differences with effect size greater than 0.2 are listed in Supplementary Table 2, with effect size calculated as the model estimate divided by the standard deviation.

For the barcode downsampling analysis, we used a custom R script to simulate downsampling 3-NICR barcodes from the 0.005%, 0.01%, and 0.015% MMS concentrations. For each tested set size (ranging from 4 to 150 3-NICR barcodes per genotype), we randomly sampled unique 3-NICR barcodes from the 0 MMS, 48-hour time point and tracked their relative abundance at the relevant MMS concentration at the same time point. A linear model was fitted to the normalized barcodes for each downsampling to detect significant differences between YYY and other genotypes and to calculate the effect size. The analysis was repeated 500 times for each tested number of 3-NICR barcodes.

Across all simulations, the frequency with which we were able to detect a statistically significant difference in growth between YYY and NNN with an effect size > 0.2 was determined. 95% confidence intervals were calculated using a binomial test through the Clopper-Pearson method.

## Data availability

Strains and plasmids are available upon request, and are described in Supplementary Tables 3 and 4, respectively. Code will be deposited on github with a permanent repository on zenodo prior to publication. Sequencing read data used in this study will be deposited at the Sequence Read Archive, and the BioProject ID will be made available prior to publication.

## Acknowledgements

We thank all members of the Sadhu lab for helpful discussions. We thank Phil Hieter for plasmids. Pacbio and Illumina sequencing was performed at the NIH Intramural Sequencing Center. This work utilized the computational resources provided by the NIH HPC Biowulf Cluster (http://hpc.nih.gov). This work was supported by the Intramural Research Program of the National Human Genome Research Institute, NIH (1ZIAHG200401).

## Supplementary material

**Supplementary table 1.** ICR barcodes were characterized as mapping to correct or incorrect gene cargos. Incorrect cargos included those that contained the wrong gene, a primer dimer, or the correct cargo with a SNP or indel. Bold indicates the ICR barcodes that were used in downstream analyses.

| Hardcoded barcode | Correct sequence | Error in primer seq. | Incorrect cargo | Primer dimer | SNP/indel |
| --- | --- | --- | --- | --- | --- |
| *RAD1* | **505** | **38** | 0 | 69 | 0 |
| *ERCC4* | **450** | **13** | 0 | 20 | 0 |
| *RAD1*-neg | **221** | **96** | 0 | 141 | 0 |
| *RAD10* | **1405** | **82** | 1 | 163 | 3 |
| *ERCC1* | **1494** | **164** | 4 | 37 | 0 |
| *RAD10*-neg | **1729** | **43** | 0 | 17 | 4 |
| *RAD14* | **709** | **19** | 1 | 33 | 0 |
| *XPA* | **1270** | **54** | 1 | 53 | 2 |
| *RAD14*-neg | **1985** | **46** | 0 | 22 | 6 |

**Supplementary table 2.** Significant differences in normalized barcode abundance for pairs of genotypes. (Included as a separate spreadsheet.)

**Supplementary table 3.** Strains used in this study. (Included as a separate spreadsheet.)

**Supplementary table 4.** Plasmids used in this study. (Included as a separate spreadsheet.)

**Supplementary table 5.** Oligonucleotides used in this study. (Included as a separate spreadsheet.)

**Supplementary figure 1.**
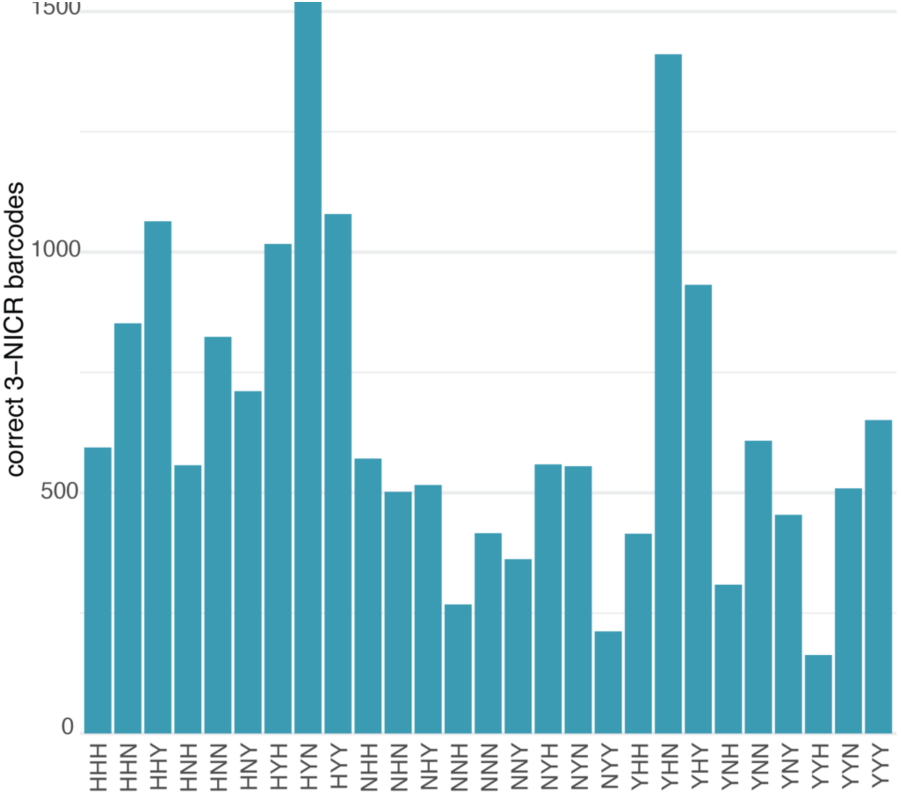
Counts of unique 3-NICR barcodes observed in the initial yeast pool. Three-letter genotype codes are as in Fig. 2c.

**Supplementary figure 2.**
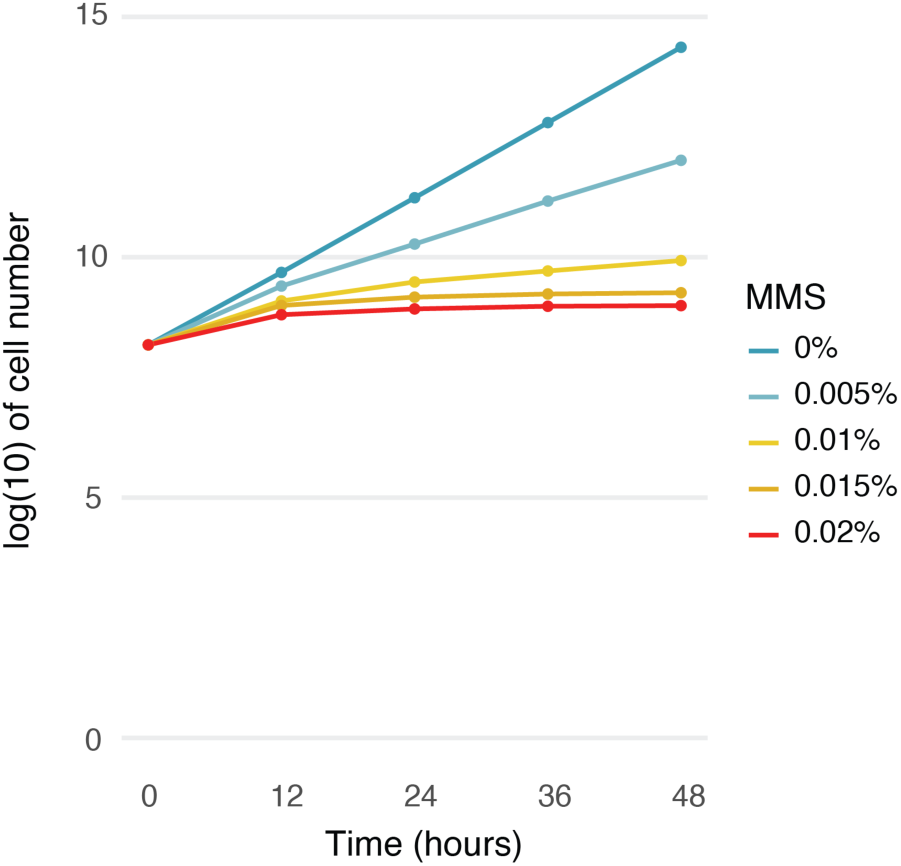
Growth in 0%, 0.005%, 0.01%, 0.015%, and 0.02% MMS of the 3-NICR yeast library cultures. Growth was measured every 12 hours as the optical density at 600 nm of each culture, with cultures that reached an optical density of 0.2 or higher diluted to an optical density of 0.025 to prevent them from reaching saturation.

## References

Adams RM, Mora T, Walczak AM, Kinney JB. 2016. Measuring the sequence-affinity landscape of antibodies with massively parallel titration curves. eLife. 5:e23156. doi:10.7554/elife.23156.

Adamson B, Norman TM, Jost M, Cho MY, Nuñez JK, Chen Y, Villalta JE, Gilbert LA, Horlbeck MA, Hein MY, et al. 2016. A Multiplexed Single-Cell CRISPR Screening Platform Enables Systematic Dissection of the Unfolded Protein Response. Cell. 167(7):1867–1882.e21. doi:10.1016/j.cell.2016.11.048.

Albert FW, Treusch S, Shockley AH, Bloom JS, Kruglyak L. 2014. Genetics of single-cell protein abundance variation in large yeast populations. Nature. 506(7489):494–497. doi:10.1038/nature12904.

Amorosi CJ, Chiasson MA, McDonald MG, Wong LH, Sitko KA, Boyle G, Kowalski JP, Rettie AE, Fowler DM, Dunham MJ. 2021. Massively parallel characterization of CYP2C9 variant enzyme activity and abundance. Am J Hum Genetics. 108(9):1735–1751. doi:10.1016/j.ajhg.2021.07.001.

Avery L, Wasserman S. 1992. Ordering gene function: the interpretation of epistasis in regulatory hierarchies. Trends Genet. 8(9):312–316. doi:10.1016/0168-9525(92)90263-4.

Becker DM, Lundblad V. 2001. Introduction of DNA into yeast cells. Curr Protoc Mol Biol. Chapter 13(1):Unit13.7. doi:10.1002/0471142727.mb1307s27.

Brideau NJ, Flores HA, Wang J, Maheshwari S, Wang X, Barbash DA. 2006. Two Dobzhansky-Muller Genes Interact to Cause Hybrid Lethality in Drosophila. Science. 314:1292–1295.

Guzder SN, Sung P, Prakash L, Prakash S. 1996. Nucleotide Excision Repair in Yeast Is Mediated by Sequential Assembly of Repair Factors and Not by a Pre-assembled Repairosome (∗). J Biol Chem. 271(15):8903–8910. doi:10.1074/jbc.271.15.8903.

Hamza A, Driessen MRM, Tammpere E, O’Neil NJ, Hieter P. 2020. Cross-species complementation of nonessential yeast genes establishes platforms for testing inhibitors of human proteins. Genetics. 214:735–747. doi:10.1534/genetics.119.302971.

Horovitz A. 1996. Double-mutant cycles: a powerful tool for analyzing protein structure and function. Fold Des. 1(6):R121–R126. doi:10.1016/s1359-0278(96)00056-9.

Johnson MS, Venkataram S, Kryazhimskiy S. 2023. Best Practices in Designing, Sequencing, and Identifying Random DNA Barcodes. Journal of Molecular Evolution. doi:10.1007/s00239-022-10083-z. https://doi.org/10.1007/s00239-022-10083-z.

Lee ME, DeLoache WC, Cervantes B, Dueber JE. 2015. A Highly Characterized Yeast Toolkit for Modular, Multipart Assembly. Acs Synth Biol. 4(9):975–986. doi:10.1021/sb500366v.

Matreyek KA, Starita LM, Stephany JJ, Martin B, Chiasson MA, Gray VE, Kircher M, Khechaduri A, Dines JN, Hause RJ, et al. 2018. Multiplex assessment of protein variant abundance by massively parallel sequencing. Nat Genet. 50(6):874–882. doi:10.1038/s41588-018-0122-z.

Nowak MA, Boerlijst MC, Cooke J, Smith JM. 1997. Evolution of genetic redundancy. Nature. 388(6638):167–171. doi:10.1038/40618.

Parvez S, Herdman C, Beerens M, Chakraborti K, Harmer ZP, Yeh J-RJ, MacRae CA, Yost HJ, Peterson RT. 2021. MIC-Drop: A platform for large-scale in vivo CRISPR screens. Science. 373(6559):1146–1151. doi:10.1126/science.abi8870.

Poelwijk FJ, Socolich M, Ranganathan R. 2019. Learning the pattern of epistasis linking genotype and phenotype in a protein. Nature Communications. 10:1–11. doi:10.1038/s41467-019-12130-8. http://dx.doi.org/10.1038/s41467-019-12130-8.

Prakash L, Prakash S. 1977. Isolation and characterization of MMS-sensitive mutants of Saccharomyces cerevisiae. Genetics. 86(1):33–55. doi:10.1093/genetics/86.1.33.

Rubin AJ, Parker KR, Satpathy AT, Qi Y, Wu B, Ong AJ, Mumbach MR, Ji AL, Kim DS, Cho SW, et al. 2019. Coupled Single-Cell CRISPR Screening and Epigenomic Profiling Reveals Causal Gene Regulatory Networks. Cell. 176(1–2):361–376.e17. doi:10.1016/j.cell.2018.11.022.

Schärer OD. 2013. Nucleotide Excision Repair in Eukaryotes. Cold Spring Harb Perspect Biol. 5(10):a012609. doi:10.1101/cshperspect.a012609.

Stevenson ZC, Moerdyk-Schauwecker MJ, Banse SA, Patel DS, Lu H, Phillips PC. 2023. High-throughput library transgenesis in Caenorhabditis elegans via Transgenic Arrays Resulting in Diversity of Integrated Sequences (TARDIS). eLife. 12:RP84831. doi:10.7554/elife.84831.

Weile J, Roth FP. 2018. Multiplexed assays of variant effects contribute to a growing genotype–phenotype atlas. Hum Genet. 137(9):665–678. doi:10.1007/s00439-018-1916-x.

Zhang J, Kobert K, Flouri T, Stamatakis A. 2014. PEAR: a fast and accurate Illumina Paired-End reAd mergeR. Bioinformatics. 30:614–620. doi:10.1093/bioinformatics/btt593. http://www.ncbi.nlm.nih.gov/pubmed/24142950.

Zhu Q, Zhao X, Zhang Y, Li Y, Liu Shang, Han J, Sun Z, Wang C, Deng D, Wang S, et al. 2023. Single cell multi-omics reveal intra-cell-line heterogeneity across human cancer cell lines. Nat Commun. 14(1):8170. doi:10.1038/s41467-023-43991-9.

